# Solution structures of the *Shewanella woodyi* H-NOX protein in the presence and absence of soluble guanylyl cyclase stimulator IWP-051

**DOI:** 10.1101/2020.08.21.262071

**Authors:** Cheng-Yu Chen, Woonghee Lee, William R. Montfort

## Abstract

Heme-nitric oxide/oxygen binding (H-NOX) domains bind gaseous ligands for signal transduction in organisms spanning prokaryotic and eukaryotic kingdoms. In the bioluminescent marine bacterium *Shewanella woodyi* (*Sw*), H-NOX proteins regulate quorum sensing and biofilm formation. In higher animals, soluble guanylyl cyclase (sGC) binds nitric oxide with an H-NOX domain to induce cyclase activity and regulate vascular tone, wound healing and memory formation. sGC also binds stimulator compounds targeting cardiovascular disease. The molecular details of stimulator binding to sGC remain obscure but involve a binding pocket near an interface between H-NOX and coiled-coil domains. Here, we report the full NMR structure for CO-ligated *Sw* H-NOX in the presence and absence of stimulator compound IWP-051, and its backbone dynamics. Non-planar heme geometry was retained using a semi-empirical quantum potential energy approach. Although IWP-051 binding is weak, a single binding conformation was found at the interface of the two H-NOX subdomains. Binding lead to rotation of the subdomains and closure of the binding pocket. Backbone dynamics for the protein are similar across both domains except for two helix-connecting loops, which display increased dynamics that are further enhanced by compound binding. Structure-based sequence analyses indicate high sequence diversity in the binding pocket, but the pocket itself appears conserved among H-NOX proteins. The largest dynamical loop lies at the interface between *Sw* H-NOX and its binding partner as well as in the interface with the coiled coil in sGC, suggesting a critical role for the loop in signal transduction.

## Introduction

H-NOX domains are gas-sensing hemoproteins that bind dissolved gases and induce signaling responses in a wide variety of organisms. In prokaryotes, H-NOX sensors function as both stand-alone proteins and as members of multidomain proteins, and respond to nitric oxide, oxygen or potentially other ligands to induce a signaling cascade [1, 2]. Obligate anaerobes use H-NOX domains to escape dioxygen as part of a methyl-accepting chemotaxis system while a variety of bacteria use stand-alone H-NOX proteins to sense nitric oxide and repress biofilm formation through lowering cyclic di-GMP levels. In the latter case, H-NOX proteins may participate in a two-component signaling cascade, inhibiting a histidine kinase that stimulates a cyclic di-GMP synthase, or by directly stimulating the phosphodiesterase activity, and inhibiting the cyclase activity, of a cyclic di-GMP synthase/phosphodiesterase fusion protein. The H-NOX protein from *Shewanella woodyi* (*Sw* H-NOX) is a member of this last category.

In animals, H-NOX domains are part of sGC proteins that generally sense nitric oxide (NO) but may also sense oxygen, for example in the worm *C. elegans* where oxygen levels regulate foraging [3, 4]. Best studied are the heterodimeric sGC proteins from insects and mammals. Heterodimeric sGC is composed of two chains, α and β, produced through gene duplication with each chain containing an N-terminal H-NOX domain, a central PAS (Per-ARNT-Sim) domain, a coiled-coil domain and a C-terminal cyclase domain [5, 6]. The α H-NOX domain has lost the ability to bind heme and is best categorized as a pseudo H-NOX domain. By producing cGMP in response to NO binding, heterodimeric sGC regulates numerous physiological processes in animals, including blood pressure, wound healing and memory formation, and is also emerging as antitumorigenic.

Loss of cGMP production through impaired sGC is linked to hypertension, atherosclerosis, asthma, neurodegeneration and contributes to heart attack and stroke. New sGC stimulating compounds, which act alone and in synergy with NO binding, are showing clinical promise and one such compound, riociguat (trade name Adempas), is in clinical use for pulmonary arterial hypertension (PAH) and related disorders [7]. sGC stimulators all derive from YC-1 (Figure 1), which was initially discovered as an antiplatelet compound [8] and eventually shown to target sGC. More recently, compound IWP-051 was developed [9], the precursor to praliciguat, which is now in clinical trial for treating hypertension in diabetics, including those with diabetic nephropathy [7].

**Figure 1.**
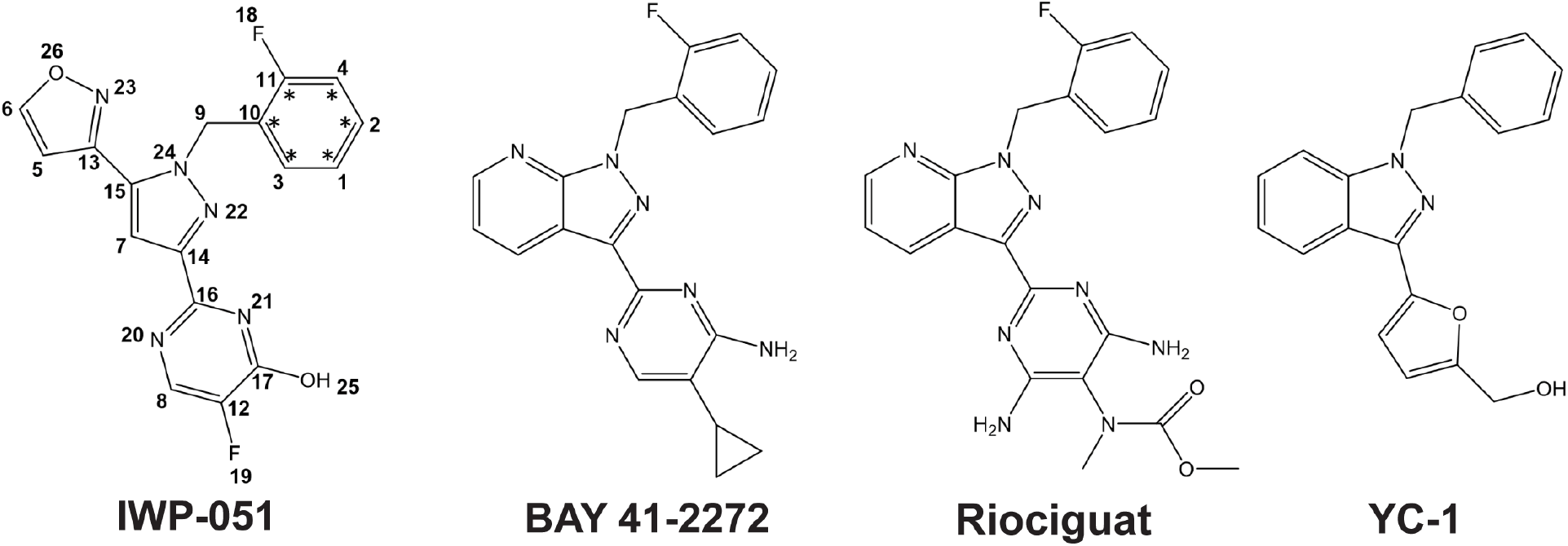
Chemical structures of sGC stimulators. The positions of ^13^C enrichment in IWP-051 is indicated with stars, and overall numbering is indicated. The 5-membered pyrazole ring is shown in the center (atoms 7, 14,15, 22 and 24) and isoxazole ring in the upper left of the figure (atoms 5, 6, 13, 23, 26).

Despite the promise of sGC stimulators, where they bind and how they work remains unclear. The H-NOX domain from *Clostridium botulinum*, also referred to as sensor of nitric oxide (*Cb* SONO), was reported to have altered CO release kinetics in the presence of sGC stimulator BAY 41-2272, suggesting binding was to the NO-sensing domain in sGC [10]. We recently found that a photoactivatable version of IWP-051 covalently attaches to residues in the β H-NOX domain and in the coiled-coil domain [11], consistent with previous studies indicating the H-NOX domain was in contact with the coiled-coil domain [12, 13]. We also showed weak binding to H-NOX proteins *Nostoc* sp. PCC 7120 (*Ns* H-NOX), *Shewanella oneidensis* (*So* H-NOX), *Shewanella woodyi* (*Sw* H-NOX) and *Cb SONO*. More recently, moderate resolution (3.8 – 5.8 Å) models for sGC in inactive and active conformations were published, showing sGC to undergo a dramatic change in conformation upon activation that involves kink-straightening and repacking of the coiled-coil domain [14, 15], which has conserved positions of instability to facilitate activation [16]. The H-NOX domain lies along the coiled coil in these structures, and in one, density for YC-1 can be seen at the interface of the H-NOX and coiled-coil domains [15].

Here, we describe the full NMR structures of *Sw* H-NOX in the absence and presence of stimulator IWP-051, which reveals a conserved pocket in the protein. We also describe the backbone dynamics for *Sw* H-NOX and how it changes in response to IWP-051 binding.

## Results

### NMR structure determination of the *Sw* H-NOX CO complex

*Sw* H-NOX was expressed in *E. coli*, purified with fully intact heme as a ferric/ferrous mixture and reduced to the ferrous state using dithionite, as previously described [11]. The protein contains the full protein (182 residues) plus eight additional residues at the C-terminus arising from the purification tag after cleavage with TEV protease. For NMR studies, we focused on the Fe(II)-CO complex, which is diamagnetic and best suited for analysis [17]. The Fe(II)-CO *Sw* H-NOX complex yielded a well-dispersed ^1^H-^15^N HSQC spectrum that was consistent with previous studies [11, 18] and unchanged for at least one week at room temperature, indicating high complex stability (Figure S1). Chemical shift assignments for *Sw* H-NOX were obtained using standard triple-resonance and NOESY experiments (described in Supplementary Information). Approximately 97% of backbone resonances and 80% of side chain resonances were assigned. The majority of unassigned resonances appear to be in the flexible loops of residues 30-45 and 81-90.

The Fe(II)-CO complex structure was determined using Xplor-NIH [19] and the following input data: NOE-derived distance restraints calibrated by NMRFAM-SPARKY [20], chemical shift-derived dihedral angle restraints from Talos-N prediction [21], hydrogen bonding restraints generated with the AUDANA algorithm [22], and global orientation restraints from residual dipolar couplings (RDCs). The AUDANA algorithm generated idealized H-bond restraints from NOE patterns and the secondary structures predicted by Talos-N. The restraints were cross evaluated with the NOESY spectrum and the structures generated during each cycle of structure calculations. Violations to the NOESY spectrum were removed during each cycle. We chose the least energy structure from 200 calculated structures for further complex structure calculation steps.

Modeling heme by NMR in heme-containing proteins is challenging due to the sparse number of NOE restraints available. To begin, we modeled *Sw* H-NOX without heme, located the heme pocket and added heme. Heme-protein NOEs guided heme placement and were consistent with the expected heme pocket from crystal structures of homologous proteins. However, the apo-model refined with a too-small heme pocket and heme-docking resulted in numerous steric clashes. To reconstitute a proper heme binding pocket, 27 loose long-range pseudo CA-CA NOE distance restraints with high lower limit boundaries were introduced for residues surrounding the heme binding pocket, which led to improved overall structure quality, improved RDC correlation (~0.98 to ~0.995) and no increase in NOE violations. These long-range restraints were removed once heme was reliably placed in the model.

Model refinement was guided by strong NOEs between the heme methine protons and protein side chains including residues L78, I79, L95, I99, I103, V107, A114, L116, A143, and L146. The Fe-NE2 distance between heme and His 104, the proximal histidine, was restrained to 2.1 ± 0.1 Å and the distance between the porphyrin ring nitrogen and His 104 NE2 restrained to 3.0 ± 0.1 Å. The Fe-CO bond was fixed at 1.9 Å and the Fe-C-O bond angle at 180°, as expected for Fe(II)-CO [23–25]. Heme methyl and vinyl groups were restrained by intra-heme NOEs derived from a 2D-[F1,F2]-^13^C^15^N-filtered NOESY experiment (Figure S2). The heme conformation was further refined with addition of heme-protein NOEs (109 in all, Table S1), which indicated a single heme orientation was present [26, 27]. NOEs were not observed for the second of the two possible heme orientations, one flipped with respect to the other, unlike in, for example, nitrophorin 4, where the two possible heme orientations exist in roughly equal proportion [27, 28]. A lack of unambiguous NOE restraints led to high flexibility in the heme propionates during docking, and steric clashes with protein loop residues 113-117, 133-138 and the N-terminal amine. H-NOX family proteins have a conserved YxSxR motif that hydrogen bonds to the propionate groups [29] and we therefore added a restraint between these residues (Y133, S135 and R137) and the propionate oxygens (5.0 Å). The N-terminal amine was additionally restrained to be near a heme propionate but with greater variability allowed (4.5 ± 0.5 Å). Including these restraints reduced steric clashes without generating NOE violations or changing backbone structure.

Heme geometry is also difficult to model by NMR, again due to the paucity of NMR restraints and the flexibility of heme. H-NOX proteins have non-planar heme geometries that vary in magnitude among the different members of the H-NOX family, adding additional complexity to modeling efforts [29–31]. To preserve heme non-planarity, we calculated topology and energy parameters optimized for non-planar heme using the semi-empirical quantum chemistry potential energy approach of MOPAC PM7, which was designed using experimental and ab initio reference data for biological systems [32]. During docking, the heme potential energy for bonds and angles was scaled to the protein potential energy (with satisfied NMR restraints), while the potential energy for heme dihedral and improper angles was left unchanged, allowing the heme to sample nonplanar conformations.

The resulting models were refined using the EEFx2 potential energy function, which simultaneously implements solvation, electrostatics, and van der Waals energy terms, allowing for a more physically realistic calculation [33]. The EEFx2 force field with implicit water was incorporated into the final structure refinement [33]. To represent the solution structure ensemble, 20 models with low energy and best data agreement (no NOE violations > 0.5 Å and no dihedral angle violations >5°) were chosen from 200 random starting models (Figure 2a). A summary of structure statistics is provided in Table S1.

**Figure 2.**
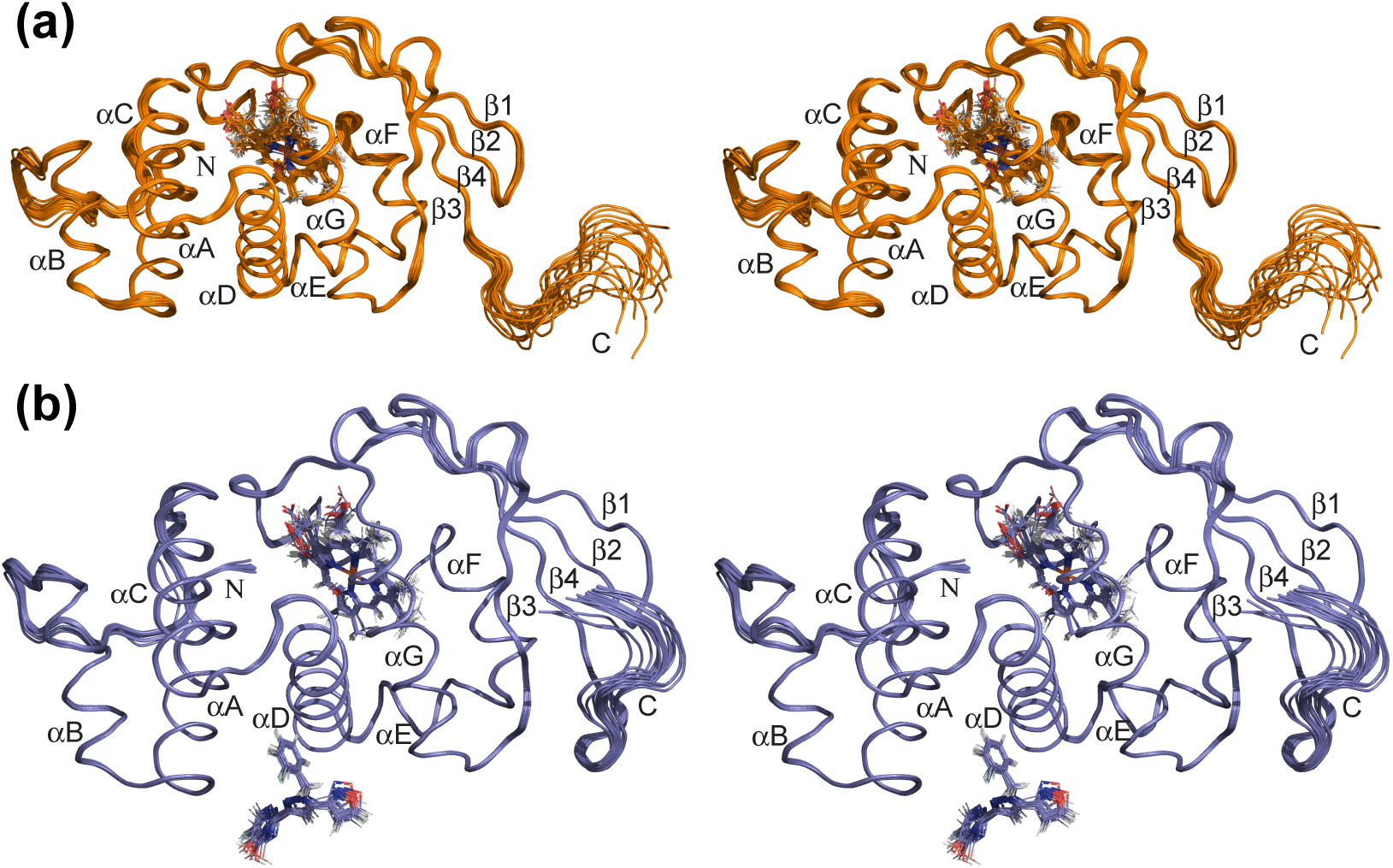
Backbone ensemble structures of *Sw* H-NOX. (a) Ligand-free protein. (b) Complex with IWP-051. Secondary structure elements are labeled and heme and IWP-051 indicated with stick representation. The structures are shown in stereo (wall-eyed view). All structure figures were prepared with PyMOL (Schrodinger, LLC).

The final *Sw* H-NOX models displayed overall fold and heme conformations that are similar to those found in other H-NOX structures, including the CO complexes from *Nostoc sp.* PCC 7120 (PDB ID 2O0G [31]), *Shewanella oneidensis* (PDB IDs 2KII [34] and 4U9G [35]), and *Caldanaerobacter subterraneus* (also known as *Thermoanaerobacter tengcongensis,* PDB ID 5JRX [36]). Key structural features include a small subdomain consisting of three helices (residues 1-61) and a large subdomain consisting of four helices, a four-stranded β sheet and the heme pocket (residues 62-182, Figure 2). The structures align remarkably well considering there is relatively low sequence identity (less than 30% in pairwise comparisons). Major differences lie mostly in the loop connecting small-domain helices B and C, large-domain β strands 3 and 4 and the packing between sub-domains.

### Structure determination of *Sw* H-NOX in complex with CO and IWP-051

Previously, we showed that stimulator compounds bind near both the coiled-coil and α H-NOX domains in sGC, and that IWP-051 interacts with bacterial H-NOX proteins in a manner similar to that for heterodimeric sGC, but with much weaker binding affinity [11]. In chemical shift perturbation studies, titration of IWP-051 to the ^15^N-labeled *Sw* H-NOX caused shifting of peaks in the ^1^H-^15^N HSQC spectrum at residues E16, F17, G18, E27, L66, L69, G71, and K73, without much linewidth broadening, indicating fast exchange binding dynamics [11]. Using transferred NOESY (TrNOESY), we found IWP-051 to occupy a single conformation on binding to bacterial H-NOX proteins and sGC [11].

To better understand binding, we determined the structure of *Sw* H-NOX bound to IWP-051 using NMR spectroscopy. *Sw* H-NOX saturated with IWP-051 yielded an ^1^H-^15^N HSQC spectrum similar to that for the protein alone and was stable during data collection (Figure S1). We included ~10% DMSO-d6 to overcome the relatively low solubility of IWP-051 in water. DMSO alone led to small shifts (~0.015 ppm) throughout the ^1^H-^15^N HSQC spectrum, while DMSO + IWP-051 led to much larger shifts [11]. In contrast, addition of phosphodiesterase inhibitor PF-04447943 in DMSO, which does not stimulate sGC but has a similar chemical composition to IWP-051, does not lead to additional shifts above the DMSO background, as previously described [11]. These data indicate IWP-051 occupies a specific binding site on *Sw* H-NOX.

To determine the structure of *Sw* H-NOX bound to IWP-051, a 5-fold excess of unlabeled compound was incubated overnight with ^13^C,^15^N-labeled *Sw* H-NOX in its Fe(II)-CO state. Data were measured for ^15^N-HSQC, ^13^C-HSQC, 2D- [F1,F2] ^13^C^15^N-filtered NOESY [37], 3D ^15^N-Edited NOESY and 3D ^13^C-Edited NOESY. A set of NH RDCs was measured with the ^15^N-labeled *Sw* H-NOX saturated by 5-fold excess of IWP-051 for dipolar coupling restraints. The majority of peaks in the HSQC and NOESY spectra were unchanged upon binding IWP-051, indicating that the protein maintained its overall native fold. We therefore used the assigned backbone resonances of unliganded *Sw* H-NOX for the dihedral restraints in the initial structure calculation. Significant shifts of NOESY peaks, due to the chemical shift perturbation on the protein amides, were found in the 3D ^15^N-Edited NOESY spectrum for the IWP-051-bound protein at residues G8, L12, E16, F17, G18, E27, L66, L69, G71, and, K73. Similar proton-proton NOESY cross peak patterns were found in these residues when compared to the unliganded protein, which allowed us to further confirm and assign the ^15^N-Edited NOESY peaks for the IWP-051 complex. The ^13^C-Edited NOESY from the IWP-051-bound protein showed relatively small chemical shift changes. The differences in the NOESY with or without IWP-051 were detected through careful inspection of the spectrum projections along the carbon or nitrogen dimension as well as spectrum segments of each residue, and entered into the refinement restraints table.

To characterize IWP-051 and heme in the complex, a 2D-[F1,F2],^13^C^15^N-filtered NOESY spectrum [37] was obtained by mixing unlabeled IWP-051 with ^13^C,^15^N-labeled *Sw* H-NOX (5:1 molar ratio, 100 ms mixing time). Heme is largely unlabeled in this sample due to the addition of unlabeled heme precursor (aminolevulinic acid) during expression. The spectrum showed similar NOESY cross peaks for IWP-051 (Figure S2) as found previously in TrNOESY spectrum [11], which indicated that a similar orientation/conformation is used by IWP-051 to bind to sGC and to the H-NOX proteins. The NOESY spectrum also showed similar intra-heme NOE cross peaks as was found in the unliganded protein (Figure S2), suggesting a similar heme environment in the IWP-051-bound protein.

The initial structure of the IWP-051-bound *Sw* H-NOX was first calculated without IWP-051 by taking into account the new NOE restraints arising from binding of IWP-051, the dihedral restraints from the native protein, the new set of NH RDCs, and the re-evaluated hydrogen bond restraints by the AUDANA algorithm. Due to weak binding of the ligand and the overcrowded NOESY spectrum, it is difficult to obtain unambiguous assignments for the Inter-IWP-051-protein NOEs from the protein NOESY spectra. To overcome this difficulty, a 2-flurobenzyl-^13^C_6_-labeled IWP-051 was synthesized and a 3D ^13^C-Edited NOESY spectrum was recorded on this compound mixed with a ^15^N-labeled *Sw* H-NOX in 100% D_2_O buffer (5:1 molar ratio, 100 ms mixing time). The ^1^H and ^13^C resonances for the 2-flurobenzyl-^13^C_6_-labeled IWP-051 can be clearly assigned using the ^1^H-^13^C HSQC, ^13^C-Edited NOESY and previously recorded 2D-COSY, 2D-TOCSY, and 2D-NOESY (Figure S2) [9, 11]. The Inter-IWP-051-protein NOEs were assigned from the 3D ^13^C-Edited NOESY (Figure S3). The initial orientation of IWP-051 was obtained by the strong NOEs from residues L12, E16, V66, and L69 to the 2-flurobenzyl ring, and the pyrazole, isoxazole and pyrimidine rings of IWP-051 were restrained by the intra-IWP-051 NOEs obtained from the 2D- [F1,F2],^13^C^15^N-filtered NOESY (Figure S2). The structure was refined in the same manner as for the unliganded protein, using Xplor-NIH [19]. 20 structures with the best agreement to the NMR restraints (no NOE violations > 0.5 Å and no dihedral angle violations > 5°) were selected from 200 random initial structures to represent the structure ensembles (Figure 2b).

The structure ensembles showed only one major conformation for IWP-051 with the 2-fluorobenzyl ring inserting into a narrow opening formed by the side chains of L12, E16 and L69 in the interface of the two sub-domains. Aside the opening is a wider hydrophobic pocket at the protein surface, formed by residues F17, V61, and L66 (Figure 3). The nitrogen atoms from pyrimidine and pyrazole ring contact the carboxylate side chains of D15 and E16, while the oxygen atom from the isoxazole ring contacts the ɛ-amino group of K72, which may lead to formation of hydrogen bonds or favorable electrostatic interactions at the binding site. This arrangement is consistent with the residues found to exhibit substantial chemical shift perturbations on binding IWP-051, which include E14, E16, F17, G18, L69, G71, and K73 [11].

**Figure 3.**
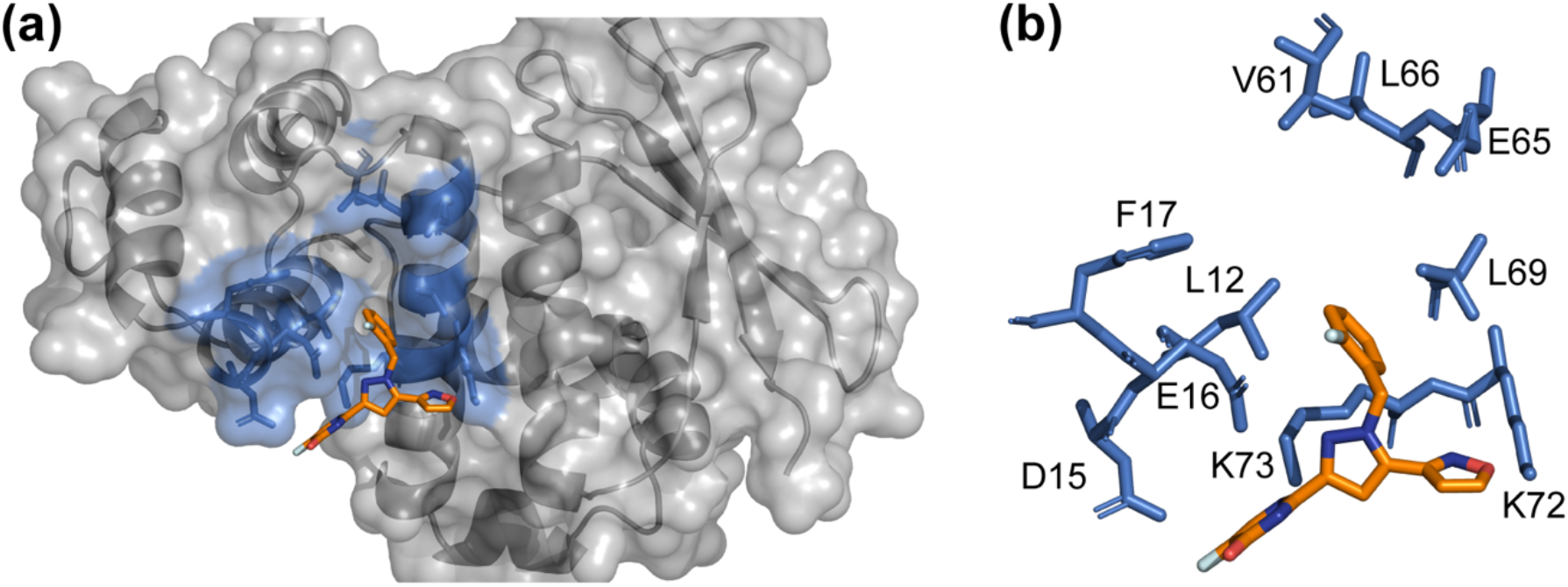
Binding interface between *Sw* H-NOX and IWP-051. (a) Surface and cartoon representation for the binding interface. Residues in the binding pocket are shown in blue. IWP-051 is colored orange and shown in stick representation. (b) Close-up view of IWP-051 binding interactions, stick representation, with IWP-051 shown in orange and binding pocket residues shown in blue. The single structure closest to the average of the ensemble structures is presented.

The IWP-051 conformation found here for binding to *Sw* H-NOX is also consistent with that expected upon binding to sGC based on TrNOESY measurements [11]. Importantly, the fluorobenzyl ring, which inserts into a crevasse between subdomains in *Sw* H-NOX, is the position in stimulator compounds least tolerant of change [9]. Compounds YC-1, BAY 41-2272 and Riociguat all retain this benzyl ring (Figure 1), as does proliciguat.

### Comparison of ligand-free and IWP-051-bound *Sw* H-NOX structures

The structure ensembles for both ligand-free and IWP-051-bound *Sw* H-NOX display good stereochemistry, internal consistency and high agreement with RDC measurements (R_dip_ = 4.8%, correlation 0.996, and R_dip_ = 2.7%, correlation 0.998, respectively; Table S1). Overall, differences between the structures are modest, generally displaying pairwise RMSD for backbone atoms of about 1 Å for the full protein, a value that drops to ~0.7 Å for the C-terminal (heme) subdomain. The smaller N-terminal subdomain displays greater variability, particularly for helix αB and the loop connecting to helix αC.

Additional analysis revealed the two subdomains in *Sw* H-NOX move toward one another on binding IWP-051, closing the binding pocket. Superposing the models using only the C-terminal subdomain (residues 62-182) highlights the change in subdomain interface, leading to backbone displacements in the small subdomain of up to 2.5 Å (Figure 4). This closed conformation leads to an improved binding pocket by bringing residues contacting each face of IWP-051 closer together (Figure 4b). Residues F17, S59 and V61, which form the upper surface of the binding pocket, also move closer to the binding site. Closure resulted in a decrease in binding pocket surface area of ~17 Å^2^ and pocket volume of ~9 Å^3^ as assessed using the CASTp algorithm [38].

**Figure 4.**
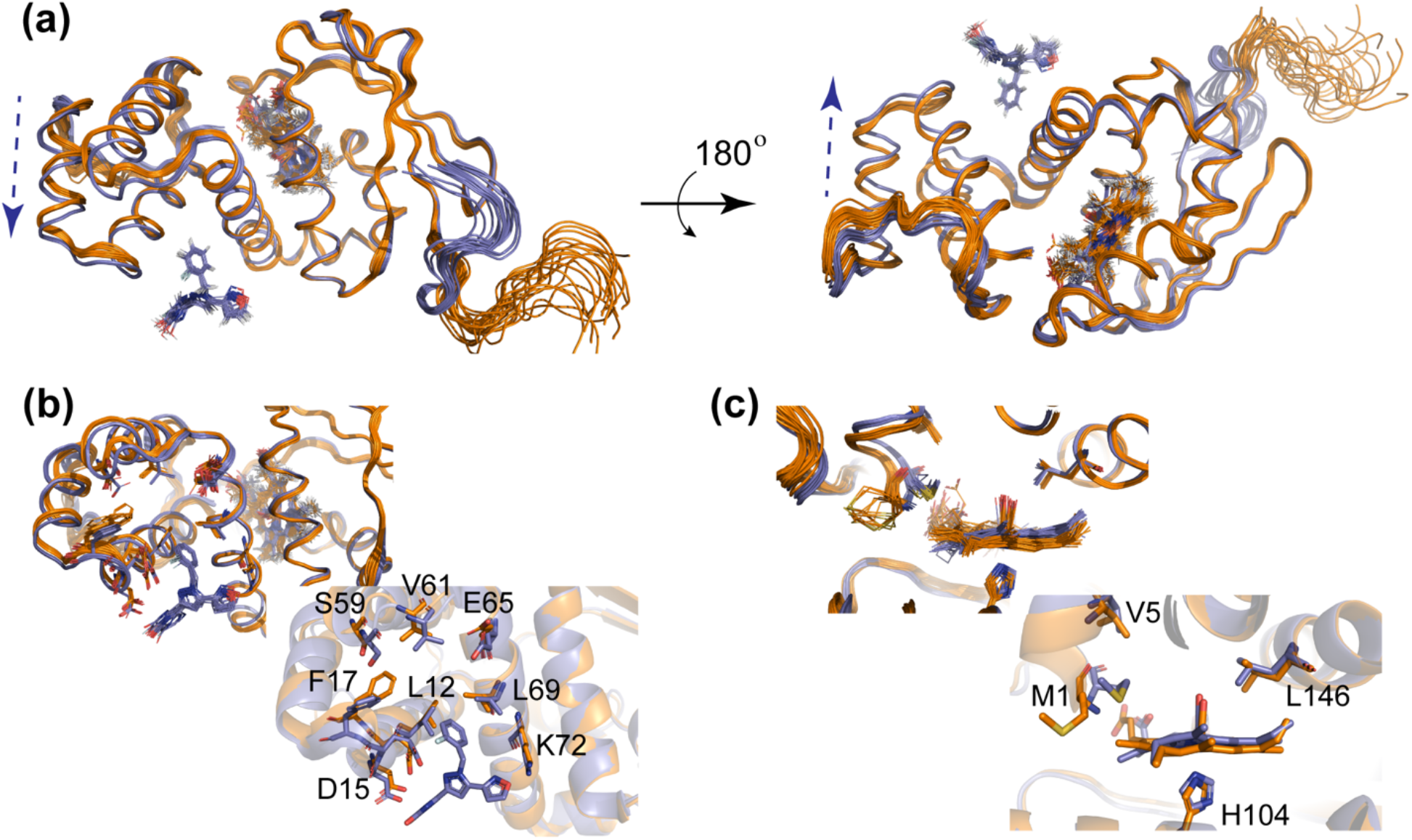
Structure comparison between unliganded and IWP-051-bound *Sw* H-NOX. (a) Unliganded and IWP-051-bound ensemble structures after superposing using only the C-terminal heme subdomain (residues 63-182). Unliganded Sw H-NOX is colored orange and the IWP-051-bound structure is colored purple. Shifting of the N-terminal subdomain on binding IWP-051 is indicated by a dashed arrow. (b) Close-up view at the IWP-051 binding site from the aligned structures (ensemble view upper left; ribbon cartoon of structures near the average structures, with labels, lower right). (c) Close-up view of the heme pocket (ensemble view upper left; ribbon cartoon, lower right).

The heme-protein NOE distance restraints allowed for well-defined heme placement in both structures. On binding IWP-051, the heme orientation is unchanged, but heme position is displaced by ~0.5 Å toward the distal heme pocket, accompanied by a similar shift in the proximal histidine (Figure 4c). Heme displacement is likely due to shifting of helix αA of the small subdomain upon IWP-051 binding. Helix αA lies on the edge of the heme distal pocket, where it contacts the distal face of heme through N-terminal residue M1. Binding pocket closure causes helix αA to translate along the distal pocked and away from the heme. Heme translation allows for the hydrophobic contact between the M1 side chain and the heme face to be maintained. On the other side of the distal pocket, L146 repacks to accommodate the new heme position. The L146 side chain contacts both the heme face and CO.

### The IWP-051 binding pocket is conserved among H-NOX proteins

We investigated whether the binding pocket uncovered in *Sw* H-NOX was unique or common among H-NOX domains. Using the CASTp algorithm [38], we searched for pockets in the H-NOX structures from *Nostoc sp* (PDB entry 2O0G [31]), *Shewanella oneidensis* (PDB entry 2KII [34]), and *Kordia algicida* (PDB entry 6BDD [36]). Surface pockets similar to that for *Sw* H-NOX were identified in all three H-NOX structures, suggesting this pocket is a conserved feature of H-NOX proteins (Figure 5a). We went on to examine binding of IWP-051 to *Ns* H-NOX (Fe(II)- CO state) using chemical shift perturbation (CSP) measurements. Titration with IWP-051 led to concentration-dependent CSPs (Figure S5), similar to those we previously observed for *Sw* H-NOX [11]. Additionally, the CSPs from *Ns* H-NOX showed fast exchange binding dynamics similar to those for *Sw* H-NOX.

**Figure 5.**
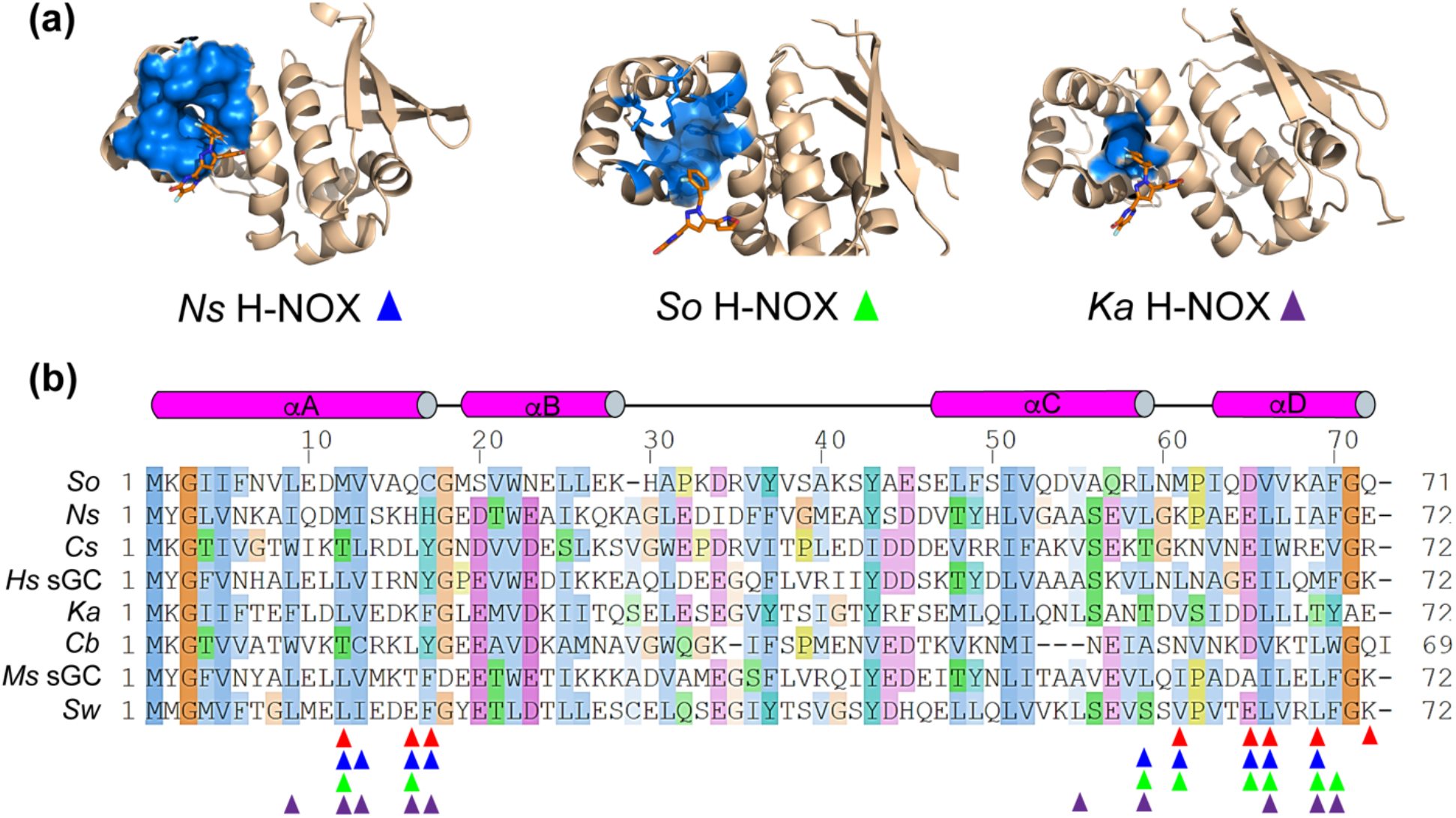
The IWP-051 binding pocket is conserved in H-NOX proteins. (a) Surface representation of the binding pockets from *Ns* H-NOX, *So* H-NOX and *Ka* H-NOX. The relative position of IWP-051 is shown by superposing the distal subdomain (residue 1-62) of *Sw* H-NOX to each structure. Residues involved in the surface pocket are shown in blue stick representation and IWP-051 in orange stick representation. (b) Multiple sequence alignment of H-NOX domains, showing binding site residues. *So*: *Shewanella oneidensis*, *Ns*: *Nostoc sp*, *Cs*: *Caldanaerobacter subterraneus*, *Hs* sGC: human sGC β H-NOX, *Ka*: *Kordia algicida*, *Cb*: *Clostridium botulinum*, *Ms* sGC: *Manduca sexta* β H-NOX, Sw: *Shewanella woodyi*. Triangles indicate the residues forming the surface pockets calculated by CASTp, with red for *Sw* H-NOX, blue for *Ns* H-NOX, green for *So* H-HNOX, and purple for *Ka* H-NOX. Secondary structure is labeled on top with α helices shown as cylinders. The alignment was performed with the program T-Coffee [48]. The graphical presentation was done by the program Jalview [49] using the Clustal X color scheme [50].

We next turned to multiple sequence alignments to search for sequence conservation in the binding pocket and extending our analysis to mammalian H-NOX domains, which do not have high-resolution structures available. Although sequence identity among these proteins is low, sequence similarity in key regions is higher. The pockets identified by CASTp generally include the same residues, indicated by triangles in Figure 5b. Of the residues that contact IWP-051 in the *Sw* H-NOX complex (indicated by red triangles in Figure 5b), sequence similarity is relatively low. We conclude that while the binding pocket is a conserved feature of H-NOX proteins, pocket chemistry is not, which would likely lead to high variability in stimulator compound affinity among members of the H-NOX family.

### IWP-051 induces a change in *Sw* H-NOX backbone dynamics

Protein conformational dynamics are central to protein function. For H-NOX proteins, binding of gaseous ligands such as NO leads to conformational changes that are key for signal transduction, whether it be to binding partners with the bacterial H-NOX proteins [18, 39], or to domain contacts leading to enhanced cyclase activity in sGC [14–16]. In sGC, binding of stimulator compounds is thought to induce similar changes to those induced on NO binding [5]. Backbone dynamical changes may play a role in H-NOX signal transduction but have not been previously reported.

To begin, we undertook NMR ^15^N-relaxation and ^15^N[^1^H] heteronuclear NOE measurements to probe the internal motion of individual residues within *Sw* H-NOX in its unliganded state. We examined the R_1_ ^15^N longitudinal relaxation rates, which are sensitive to fast (picosecond to nanosecond) backbone motions [40], and found uniform backbone dynamics throughout the protein except for increased flexibility in two loop regions, spanning residues 28-45 and 88-91 (Figure 6a). The R_2_ ^15^N transverse relaxation rates, which not only respond to fast (nanosecond) motions but also to slower (microsecond to millisecond) chemical exchange processes, were also uniform throughout the protein except for residues in loop 28-45. In this loop, R_2_ values nearly doubled for residues Q32, V40, and H45 (Figure 6b). Finally, we measured ^15^N[^1^H] heteronuclear NOE, which provide information about individual backbone dynamics and is sensitive to fast (picosecond) backbone motions. Residues undergoing faster dynamics show a decrease in NOE intensity, measured as a ratio of peak intensities with and without saturation. Here again, residues 28-45 displayed increased backbone dynamics (Figure 6c). Importantly, on examining structures of H-NOX proteins determined by X-ray crystallography, the backbone atoms for residues 28-45 are generally well-determined and do not display high disorder with the exception of residue 32, which in some cases is poorly ordered. Sidechains for this region often display disorder in crystal structures and the backbone conformations for this stretch vary widely, suggesting crystal contacts may stabilize one of several conformations for this region of the protein. Here, using NMR structure determination (Figure 2) and dynamics measurements (Figure 6), we observe higher variability in residues 28-45.

**Figure 6.**
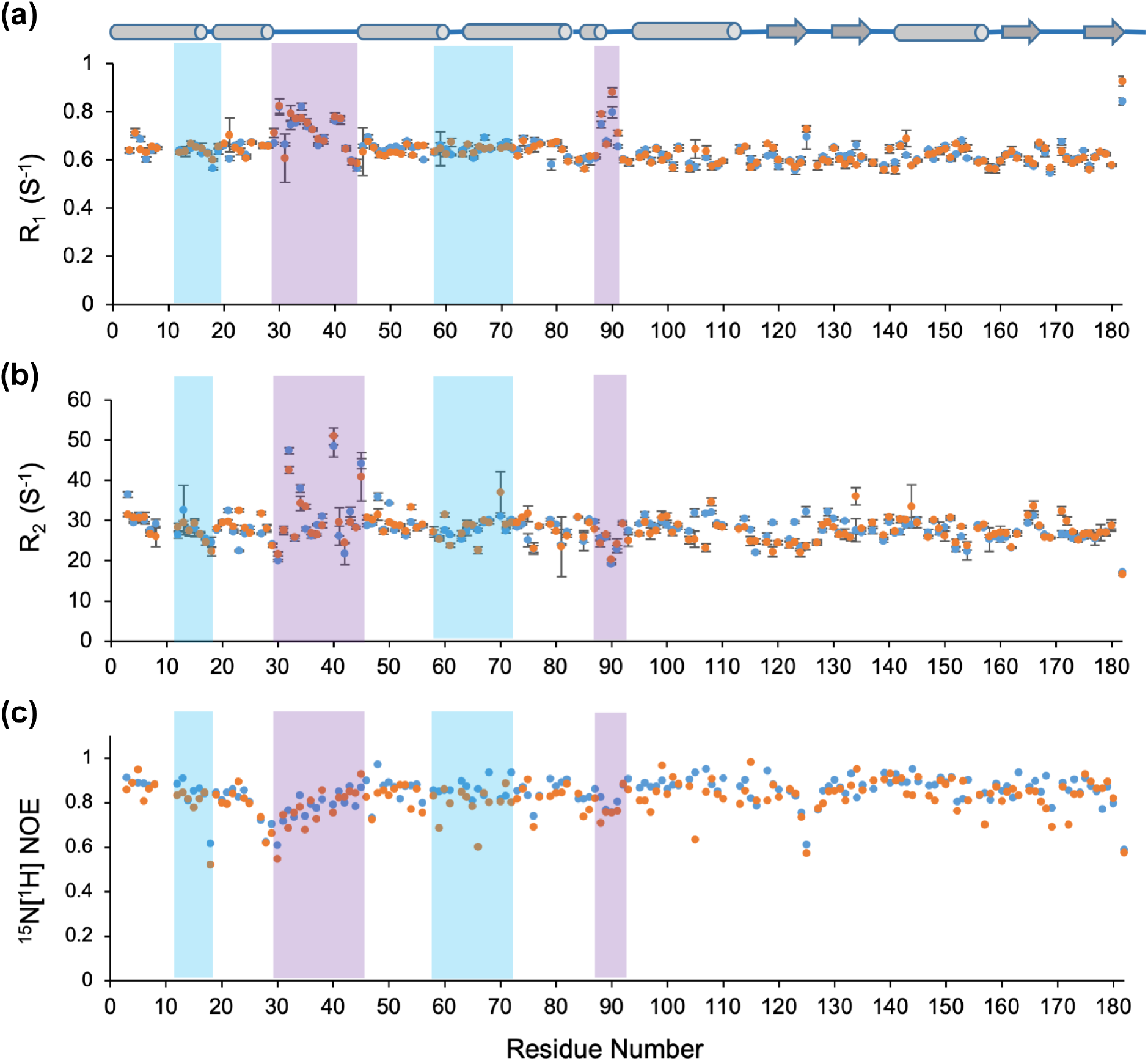
^15^N relaxation dynamics for native and IWP-051-bound *Sw* H-NOX. (a) ^15^N longitudinal relaxation rates (R_1_) plotted against residue number. (b) ^15^N transverse relaxation rates (R_2_) plotted against residue number. (c) ^15^N[^1^H] heteronuclear NOE intensities plotted against residue number. Secondary structure elements are indicated by cylinders for α helices and arrows for β strands. Data points for unliganded and IWP-051-bound proteins are shown as blue and orange dots, respectively. Error bars are the standard deviation from data fitting (R_1_ and R_2_ plots). Flexible loops are highlighted with purple boxes and the IWP-051 binding site is highlighted with blue boxes.

Binding of IWP-051 did not greatly alter *Sw* H-NOX dynamics. A similar R_1_ distribution was observed with the exception that the loop region 88-91 gained more flexibility (Figure 6a), a region well-away from the binding site. In R2 measurements, loop 28-45 displayed somewhat dampened dynamics, particularly for residues Q32, E34 and H45 (Figure 6b). HetNOE values (Figure 6c) decreased at the binding site, including those for G18, S59, V61, and L66, indicating IWP-051 binding leads to greater backbone motion at the binding site. Interestingly, similar residues, including E16, F17, and E20, are involved in *Sw* H-NOX recognition of its binding partner, cyclic-di-GMP synthase [18]. Flexibility in this region may facilitate binding.

### Comparison with photoaffinity labeling and cryo-EM studies

Although high resolution structural data for stimulator binding to sGC are still lacking, insight into binding location come from photoaffinity covalent labeling combined with mass spectrometry [11], and from single-particle cryo-electron microscopy [15]. Photolabeling was conducted with a stimulator compound modified to have a diazirine moiety attached to the pyrimidine ring, where modifications are well tolerated, and photoactivation leads to covalent attachment to nearby residues in the protein. For sGC from *Manduca sexta*, these data indicate that stimulators bind near both the β H-NOX domain (residues 6-9) and the coiled-coil domain (β chain residues 361-362 and 365-366). A recent cryo-EM study of sGC bound to YC-1, also using the protein from *Manduca sexta*, yielded a backbone model of sGC at 5.8 Å resolution with density attributed to YC-1 in two possible orientations. In this model, YC-1 was within 5 Å of β H-NOX residues V39, F77, C78, and Y112, as well as coiled-coil residues β Q349 and α 422-427.

Our position for IWP-051 in *Sw* H-NOX is consistent with both models, although not perfectly overlapping. In particular, four of five of the residues either labeled through photoaffinity or identified by cryo-EM in the H-NOX domain, lie along the same subdomain interface identified for binding in the NMR structure, and the fifth residue is nearby. However, the binding site found for *Sw* H-NOX is translated a few angstroms along the interface, moving away from the heme distal pocket, with respect to the cryo-EM and photoaffinity labeling studies.

## Discussion

We have determined the NMR structure of *Sw* H-NOX in the presence and absence of an sGC stimulator, IWP-051, to high precision. Spectra were measured with heme in the ferrous (Fe II) state and bound to CO, yielding a diamagnetic system amenable to full structure determination. The protein displays the expected overall H-NOX fold, with a small N-terminal helical domain and a lager C-terminal domain of mixed α helix and β sheet and a large heme pocket. The heme conformation is modestly distorted from planar and is linked to the protein through a proximal histidine, H104. The distal pocket is hydrophobic and with extra room, typical of H-NOX proteins. The sidechain for L146, however, is in contact with both CO and heme.

Stimulator IWP-051 binds only weakly to *Sw* H-NOX (*K*_d_ ~1 mM), unlike for sGC where binding is tighter and stimulates sGC in HEK cells with an EC_50_ value of 290 nM [9]. Binding to *Sw* H-NOX is, however, specific and involves the same compound conformation as for binding to sGC [11], with the benzyl and pyrimidine rings perpendicular to the core pyrazole ring, and the isoxazole ring rotated slightly out-of-plane with respect to the pyrazole ring. The benzyl ring, which is the position least tolerant to change in sGC stimulators (Figure 1; [7]), inserts between the large and small subdomains, contacting residues L12, E16 and L69. The pyrazole ring is also near E16 and in an arrangement that could lead to hydrogen bond formation if E16 were protonated. The isoxazole and pyrimidine rings are less in contact with the protein, but are close to K72 and D15, respectively. Overall, binding interactions are modest, consistent with the weak affinity of IWP-051 for *Sw* H-NOX. Binding does however induce closure of the binding pocket through an ~2.5 Å shift in small-subdomain position with respect to large subdomain. This rotation is similar to that observed in the H-NOX from *Shewanella oneidensis* on binding NO [1, 35].

In addition to subdomain movements, dynamical properties of H-NOX backbone residues are likely of importance in forming productive complexes with binding partners, and in signal transduction in multidomain proteins such as sGC. Here, we show that *Sw* H-NOX has relatively uniform backbone dynamics across both subdomains with the exception of the large loop between helices αB and αC (residues 28-45, small subdomain) and the loop between helices αE and αF (residues 88-91, large subdomain), both of which show enhanced dynamics compared to the overall protein. Dynamics in these regions increase on binding IWP-051, as do dynamics in the binding site. Interestingly, residues 38 and 40 directly contact the sGC coiled coil in the activated conformation observed by cryo-EM, but not in the inactive form [14, 15].

How stimulator binding to *Sw* H-NOX relates to binding in full-length sGC remains unknown, but certain aspects are becoming clear. Importantly, both photochemical labeling [11] and cryo-EM [15] support a model in which stimulator compounds bind at the interface of the H-NOX and coiled-coil domains, and therefore stand-alone H-NOX domains contain only a portion of the binding site. Nonetheless, the binding pocket identified in *Sw* H-NOX appears to be conserved among H-NOX proteins (Figure 5). Binding to the subdomain interface, as we report here for *Sw* H-NOX, appears to also occur in sGC, although possibly to a position somewhat shifted toward the heme distal pocket. Additionally, movement at the subdomain interface as occurs for *Sw* H-NOX upon binding IWP-051 is also seen on activation of sGC when examined by cryo-EM [14]. Interestingly, mutations introduced into the subdomain interface can either block stimulator binding, as seen with the L12W/T48W double mutant in *Manduca* sGC [11], or mimic stimulator action, as seen in the I52W/L67W double mutant, also in *Manduca* sGC*.

## Materials and methods

Detailed methods are described in Supporting Information.

### Protein expression, purification, and NMR sample preparation

Proteins were expressed in *E. coli* and purified using a His-tag fused to the protein followed by cleavage of the tag with TEV protease, as previously described [11]. Isotope enrichment was accomplished using M9 media isotopically enriched for ^15^N or ^13^C/^15^N. Full protocols for protein isolation and preparation for NMR experiments are described in the Supplemental information text.

### NMR spectroscopy

NMR experiments were performed at 293 K on either Agilent 800-MHz or Bruker 600, 800-MHz Avance NEO spectrometers, all equipped with triple resonance cryogenic probes. Samples for NMR measurements typically contained 0.5-0.8 mM protein in buffer containing 90% H_2_O/10% D_2_O or 100% D_2_O, and 50 mM Na/KPO_4_, 50 mM NaCl at pH 7.4. The Fe(II)-CO complex was prepared by reducing heme with dithionite followed by saturation with CO, as previously described [11] and in the Supplemental Information text. IWP-051 was dissolved in 100% DMSO-d6 and 25-50 mM was prepared as a stock solution. For the *Sw* H-NOX IWP-051 complex, a 5-fold excess amount of IWP-051 was used in the NMR measurements, which resulted in ~10-15% DMSO-d6. Both labeled and unlabeled IWP-051 were generously provided by Ironwood Pharmaceuticals.

### NOE distance restraints, dihedral angle, hydrogen bond, and RDC restraints

Intramolecular NOE distance restraints for protein were obtained from 3D ^15^N-NOESY-HSQC, 3D ^13^C-HSQC-NOESY (in 100% D_2_O buffer) and 3D ^13^C-HSQC-NOESY (aromatic). TALOS-N was used to predict backbone dihedral angles from chemical shifts [21]. Hydrogen bond restraints were generated using the AUDANA algorithm, which predicts hydrogen bonds from NOE cross peak patterns of secondary structures and reevaluates them after each cycle of structure calculation [22]. RDC data were obtained using 2D ^1^H-^15^N correlation via TROSY allowing for TROSY and AntiTROSY signal in an IPAP manner [41]. The RDC constants were obtain by the differences between TROSY and AntiTROSY components from unaligned and pf1 phage-aligned samples.

### Structure determination

NOESY peaks were assigned manually and automatically in iterative cycles of PONDEROSA-C/S’s AUDANA and CYANA calculations [42, 43]. Peak intensities were calibrated and converted to distance restraints using NMRFAM-SPARKY [20]. Structure determination was accomplished using Xplor-NIH [19]. The PDB, topology and parameter files for heme and IWP-051 were prepared using Open Babel [44] and RUNER [45], and using MOPAC PM7 force fields with General Amber Force Field (GAFF) options [32]. The MOPAC PM7 force field was designed using experimental and ab initio reference data for biological systems. Open Babel is a tool for format conversion. RUNER was the front end developed by the National Magnetic Resonance Facility at Madison (NMRFAM) to unify the nomenclatures using the ALATIS tool for labeling consistency [46]. PSF files were generated by Xplor-NIH. The detailed structure calculation is described in the Supplemental Information text.

### Chemical shift perturbation for *Ns* H-NOX

CSPs were measured by collecting a series of ^1^H-^15^N HSQC spectrum using 450 μM ^15^N-labeled *Ns* H-NOX with increasing concentration of IWP-051 (100-4000 μM) or DMSO-d6 (1.6-9.6%). Peaks with significant chemical shift change were identified by overlaying the spectra. The shifts due to the effects from DMSO-d6 were ignored.

### ^15^N relaxation and ^15^N[^1^H]-heteronuclear NOE (hetNOE) measurements

The ^15^N longitudinal (R_1_) and transverse (R_2_) relaxation rates, and ^15^N[^1^H] heteronuclear NOE, were measured in an interleaved manner as in the TROSY-based experiments [47] at 800 MHz. A recycle delay of 3.0 s between experiments was used along with the following relaxation delays for T1: 50, 200, 400, 800, 1200, 1600, 2000, 2500, 3000 ms; and for T2: 8.5, 17.0, 25.5, 33.9, 42.4, 50.9, 67.87, 84.84, 101.81, and 118.8 ms. The spectra were analyzed and plotted by NMRFAM-Sparky to obtain the rates for each residue. The ^15^N[^1^H]-NOE was determined from a pair of interleaved spectra acquired with or without proton presaturation at 800 MHz. Values of hetNOE were obtained by ratio of the peak intensities for each residue.

### Accession numbers

Coordinates for the *Sw* H-NOX structures are deposited in the Protein Data Bank under PDB entries 6OCV and 6WQE. NMR assignment data are deposited in the Biological Magnetic Resonance Data Bank under BMRB entry 27284.

## Abbreviations used

AUDANA: Automated Database-Assisted NOE Assignment
BMRB: Biological Magnetic Resonance Bank
cryo-EM: cryogenic electron microscopy
CSP: chemical shift perturbation
*Cb*: *Clostridium botulinum*
*Cs*: *Caldanaerobacter subterraneus*
hetNOE: heteronuclear NOE
H-NOX: heme-nitric oxide/oxygen binding
HSQC: heteronuclear single quantum coherence
*Ka*: *Kordia algicida*
*Ms*: *Manduca sexta*
PDB: Protein Data Bank
PONDEROSA: Peak-picking Of NOE Data Enabled by Restriction of Shift Assignments
NMR: Nuclear Magnetic Resonance
NMRFAM: National Magnetic Resonance Facility at Madison
NO: nitric oxide
NOE: nuclear Overhauser effect
NOESY: nuclear Overhauser enhancement spectroscopy
*Ns*: *Nostoc* sp. PCC 7120
PAH: pulmonary arterial hypertension
PAS: Per-ARNT-Sim
RDC: residual dipolar coupling
RMSD: root-mean-square deviation
sGC: soluble guanylyl cyclase
*So*: *Shewanella oneidensi*
TrNOESY: transferred NOESY
TROSY: transverse relaxation optimized spectroscopy

## Supplemental data

Supplementary text and data to this article can be found online at: XXX.

## Acknowledgments

We are grateful to Michael Clarkson for help with NMR measurements at the University of Arizona, and to Ironwood Pharmaceuticals for providing labeled and unlabeled IWP-051. This study was supported by National Institutes of Health Grants R01 GM117357, P30 CA023074 and U54 CA143924 (to W. R. M.) and T32 HL007249 (to C. C.). This study was also supported by Grants 17POST33670593 (to C. C.) from the American Heart Association and by Sponsored Research Agreement 100003104 from Ironwood Pharmaceuticals (to W. R. M.). This study made use of the National Magnetic Resonance Facility at Madison, which is supported by NIH grants P41 GM103399 (NIGMS) and P41 GM66326 (NIGMS). Additional equipment was purchased with funds from the University of Wisconsin, the NIH (RR02781, RR08438), the NSF (DMB-8415048, OIA-9977486, BIR-9214394), and the USDA.

## Footnotes

Double mutation (I52W/L67W) in the heme domain of soluble guanylyl cyclase mimics stimulator compounds, Jessica A. Kievenaar, Andrzej Weichsel, Jacob T. Croft, Jinghui Li, Changjian Feng, and William R. Montfort, *manuscript in preparation*.

## Author Contributions

**Cheng-Yu Chen:** Conceptualization, Investigation, Formal analysis, Methodology, Funding acquisition, Writing-Original draft preparation, Reviewing and Editing. **Woonghee Lee:** Investigation, Formal analysis, Methodology, Writing-Reviewing and Editing. **William R. Montfort:** Conceptualization, Project administration, Funding acquisition, Formal analysis, Writing-Original draft preparation, Reviewing and Editing.

