## Supplemental Information for "Solution structures of the *Shewanella woodyi* H-NOX protein in the presence and absence of soluble guanylyl cyclase stimulator IWP-051"

| | Supplementary Results | Page |
|---|---|---|
| **Suppl. Text** | Methods | 3 |
| **Figure S1** | $^{1}H$-$^{15}N$ HSQC spectra overlay of ligand-free and IWP-051-bound *Sw* H-NOX | 7 |
| **Figure S2** | Assignments for protein-bound heme and IWP-051 | 8 |
| **Figure S3** | Segments from a 3D $^{13}C$-NOESY-HSQC measured on $^{13}C$-benezyl-labeled IWP-051 bound to $^{15}N$-*Sw* H-NOX | 9 |
| **Figure S4** | Multiple sequence alignment of H-NOX domains | 10 |
| **Figure S5** | Superimposed $^{1}H$-$^{15}N$ HSQC spectra of *Ns* H-NOX titrated with IWP-051. | 11 |
| **Table S1** | Structure statistics | 12 |
| **Supp. Ref.** | | 13 |

### Supplementary Information Text

#### Protein expression and purification

The genes encoding the H-NOX domains from *Shewanella woodyi* (*Sw* H-NOX) and *Nostoc* sp. PCC 7120 (*Ns* H-NOX) were synthesized and cloned into pET21b+ for expression (GeneScript Biotech Corp). *Sw* H-NOX was expressed in *E. coli* Tuner (DE3) pLysS cells. Cultures were grown at 37 °C in M9 media isotopically enriched for $^{15}N$ or $^{13}C/^{15}N$ until $OD_{600}$ reached ~0.8 and then the temperature was cooled down to 20 °C. Expression was induced by addition of 0.5 mM isopropyl β-D-thiogalactopyranoside (IPTG) and supplemented with 0.5 mM δ-aminolevulinic acid (ALA) to improve heme synthesis. *Ns* H-NOX was expressed in *E. coli* Rosetta cells. The expression was induced by 0.25 mM IPTG and 0.25 mM ALA and the induced cells were grown at 20 °C for an addition of 20 hr. Purification of *Sw* and *Ns* H-NOX was carried out as previously described [1]. All buffers were extensively degassed prior to usage. Briefly, cells were lysed by French press in buffer A (50 mM Tris-HCl, pH 8.0, 250 mM NaCl, 1.0 mM TCEP and 1.0 mM PMSF). The proteins were purified using a HisTrap FF nickel-nitrilotriacetic acid affinity column (GE Healthcare) with buffer A supplemented with 30-60 mM EDTA, followed by TEV cleavage. The final purification was performed by size exclusion chromatography using a superdex 75 column (GE Healthcare) pre-equilibrated in 20 mM Tris-HCl (pH 8.0), 100 mM NaCl and 5 mM β-mercaptoethanol. Protein concentrations were measured using the Pierce 660-nm protein assay kit (Thermo Fisher Scientific) and samples were frozen in liquid nitrogen for storage at -80 °C.

#### NMR sample preparation

Both *Sw* and *Ns* H-NOX were prepared in their Fe(II)-CO diamagnetic states as described previously [1]. Briefly, 10 mM dithionite and 2 mM dithionite were used to reduce *Sw* H-NOX and *Ns* H-NOX, respectively, in an extensively degassed buffer containing 50 mM Tris-HCl (pH 8.0) and 50 mM NaCl, and then saturated with CO in a sealed tube. Samples were then extensively buffer-exchanged to the NMR buffer containing 50 mM sodium/potassium phosphate (pH 7.4), 50 mM NaCl, 10% $D_2O$ and saturated CO. NMR experiments were performed in a CO-saturated and sealed shigemi tube. The same NMR buffer but in 100% $D_2O$ was used for the 3D $^{13}C$-HSQC-NOESY, 3D $^{13}C$-NOESY-HSQC and, 2D $^{13}C^{15}N$-filtered [F1,F2] NOESY experiments. The concentration of NMR samples were 0.5-0.8 mM. An $^{15}N$-HSQC spectrum was collected after each experiment to evaluate the stability of the protein. IWP-051 was prepared by dissolving the compound into DMSO-d6 (99.9 atom % D, Aldrich) to a final concentration of 25-50 mM as stock solution. 2,2-dimethyl-2-silapentane-5-sulfonate (DSS) was included as an internal reference. Samples for residual dipolar coupling (RDC) measurements were prepared by the addition of pf1 phage (Asla Biotech) to a final concentration of ~15 mg/mL [2]. The $D_2O$ quadrupolar splitting was measured before and after each experiment to validate the alignment.

#### NMR Spectroscopy

NMR experiments were carried out on Agilent 18.8 T (800 MHz) NMR spectrometer or on Bruker 600 and 800 Mhz spectrometers, all equipped with a triple resonance cryogenic probe. All experiments were performed at 20 °C. NMR data were processed using NMRPipe [3] and in combination with MddNMR [4, 5] or SMILE [6] for non-uniformly sampled data. NMRFAM-SPARKY [7], I-PINE [8], and PINE-SPARKY [9, 10] were used for resonance assignment and spectrum analyses. The program TALOS-N was used to estimate backbone dihedral angels from chemical shifts [11].

**Assignment of protein backbone, side chain, heme and IWP-051 resonances**

All protein backbone and side-chain resonance experiments were acquired using 40-50% non-uniform sampling with schedules generated by the Poisson Gap Sampling Method [12]. The sequence-specific backbone resonances were assigned using the following triple resonance experiments: 3D-HNCA [13], 3D-HN(CO)CA, 3D-HNCACB [13], 3D-CBCA(CO)NH [14], 3D-HNCO [15, 16], and 3D-(HCA)CO(CA)NH [17]. Side-chain resonance assignments were obtained using 3D-HBHA(CO)NH [18], 3D-C(CO)NH [19], 3D-H(CCO)NH [19], 3D-HCCH-TOCSY [20], and 3D-HCCH-COSY [20]. The aromatic side chains were assigned using 2D constant time $^{1}H$-$^{13}C$ HSQC and 3D $^{13}C$-HSQC-NOESY (100-ms mixing time) [21]. The $^{1}H$ resonances for the protein-bound HEME were assigned using 2D $^{13}C$-filtered [F1,F2] NOESY (100-ms mixing time) [22]. The $^{1}H$ resonances for the free IWP-051 were assigned using 2D-COSY, 2D-TOCSY, 2D-NOESY as described previously [1, 23]. The $^{1}H$ resonances for protein-bound IWP-051 were assigned by 2D $^{13}C$-filtered [F1,F2] NOESY [22], and 2D-Transffered-NOESY [1]. The $^{13}C$ resonances for protein-bound IWP-051 were assigned by $^{1}H$-$^{13}C$ HSQC and 2D $^{13}C$-filtered [F1,F2] NOESY.

**Distance restraints**

Intramolecular distance restraints for native and the IWP-051-bound *Sw* H-NOX were obtain from 3D $^{15}N$-NOESY-HSQC [24] and 3D $^{13}C$-HSQC-NOESY (in 100% $D_2O$ buffer) [21], both collected with 100-ms mixing time. Intermolecular distance restraints between protein and heme were obtained from 3D $^{13}C^{15}N$-filtered [F1,F2] NOESY-HSQC and 3D $^{13}C$-HSQC-NOESY in 100% $D_2O$ buffer, both collected with 100-ms mixing times and in 100% D2O. Intramolecular distance restraints for heme and IWP-051 were obtained from 2D $^{13}C$-filtered [F1] NOESY with 100-ms mixing times. Intermolecular distance restraints between protein and IWP-051 were obtained from 3D $^{13}C$-NOESY-HSQC [24] on a $^{12}C$ protein and specifically $^{13}C$-enriched IWP-051 sample. Hydrogen bond restraints for the protein were obtained from AUDANA algorithm which analyzes NOE patterns with TALOS-N secondary structure predictions, and eliminates violations from each cycle of structure calculation [25].

**Residual dipolar coupling measurements**

RDC data were obtained at 800 MHz using 2D $^{1}H$-$^{15}N$ correlation via TROSY allowing for TROSY and AntiTROSY signal in an IPAP manner [26]. The RDC constants were measured by the differences between TROSY and AntiTROSY components from unaligned and pf1 phage aligned samples.

**Structure calculation**

A heme-free apo structure was first calculated using the NOE distance restraints, backbone dihedral restraints, hydrogen bond restraints and RDCs. Initial structures were obtained through iterative cycles of manual NOE assignment followed by cycles of automated NOE assignment with both CYANA and AUDANA algorithms in the PONDEROSA-C/S software package [25, 27, 28]. The structures were further refined by Ponderosa Refinement-X option using simulated annealing and the molecular dynamics protocol in Xplor-NIH [29] along with RDCs and with additional restraints generated subsequently by AUDANA [25]. NOE restraints were manually validated using the Distance Constraint Validator in the Ponderosa Analyzer which visualizes and validates the NOE peaks with the corresponding restraint distances in the calculated structures and NOESY spectrum [28]. The backbone dihedral restraints were derived from chemical shifts using TALOS-N [11].

The structure PDB, topology and parameter files for heme and IWP-051 were prepared using Open Babel [30] and RUNER [31] and with force fields MOPAC PM7 with General Amber Force Field (GAFF) [32] options. PSF files were generated by Xplor-NIH [29]. Heme was incorporated into the protein using additional inter- and intra-molecular NOE distance restraints along with restraints from protein to obtain an initial heme position and orientation. The potential energy of bond and angle in heme was scaled to 0.5 of the potential energy of protein defined in Xplor-NIH. The complex was further refined by the molecular dynamics routine in Xplor-NIH. The molecular dynamic routine includes 3500 K high temperature dynamics and the thermal bath was slowly cooled to 25 K along with IVM rigid body modeling, followed by low temperature torsion angle dynamics, and cartesian dynamics. A total of 200 structures were calculated using this protocol. No NOEs could be identified between the heme propionates and the protein, which resulted in steric clashes between the propionate sidechains and the protein. Crystallographic and sequence data have shown that a conserved YxSxR motif in the H-NOX family of proteins forms hydrogen bond interactions to the heme propionate [33, 34]. Based on this assumption and with a conserved YxSxR motif found in *Sw* H-NOX, a loose five Å distance restraint between side-chain protons of Y133, S135, R137 and the propionate motif were introduced to restrain the positions of the propionate atoms. The propionate was further restrained addition of hydrogen bond restraints from the N-terminal amine to the propionate carboxylate, as found in H-NOX crystal structures [33], to avoid steric clashes. Addition of these restraints did not result in the violation of any experimental distance restraints or changing of the protein backbone structure. Four to seven cycles of iteration were conducted, with NOE violations removed or adjusted in each cycle. At the final stage of refinement, an implicit solvation potential, EEFx2 was used to refine the structures [35]. A total of 200 models were calculated and refined against the distance restraints, dihedral angle restraints, hydrogen bond restraints and RDCs using the molecular dynamics routine with the EEFx2 potential. The 20 models with lowest energy and no NOE violations > 0.5 Å were selected to represent the final ensemble for both ligand-free and IWP-051-bound *Sw* H-NOX structures. The quality of structure was evaluated with the protein structure validation software suite (PSVS) [36].

#### $^{15}N$ relaxation and $^{15}N[^1H]$-NOE (hetNOE) measurements

The $^{15}N$ longitudinal ($R_1$) and transverse ($R_2$) relaxation rates and $^{15}N[^1H]$ heteronuclear NOE were measured in the same manner as in the TROSY-based experiments [37] at 800 MHz. A recycle delay of 3.0 s between experiments and the following relaxation delays were used for T1: 50, 200, 400, 800, 1200, 1600, 2000, 2500, 3000 ms; and for T2: 8.5, 17.0, 25.5, 33.9, 42.4, 50.9, 67.87, 84.84, 101.81, and 118.8 ms. The spectra were analyzed and plotted by the *Peak Intensity Analysis* plugin tool (two-letter-code *rh*) in NMRFAM-Sparky to obtain the relaxation rates for each residue. The effects due to DMSO (~10%) were normalized using the average rates from core secondary structures. The $^{15}N[^1H]$-NOE measurements were determined from a pair of interleaved spectra acquired with or without presaturation of proton at 800 MHz. Values of hetNOE was obtained by ratio of peak intensities.

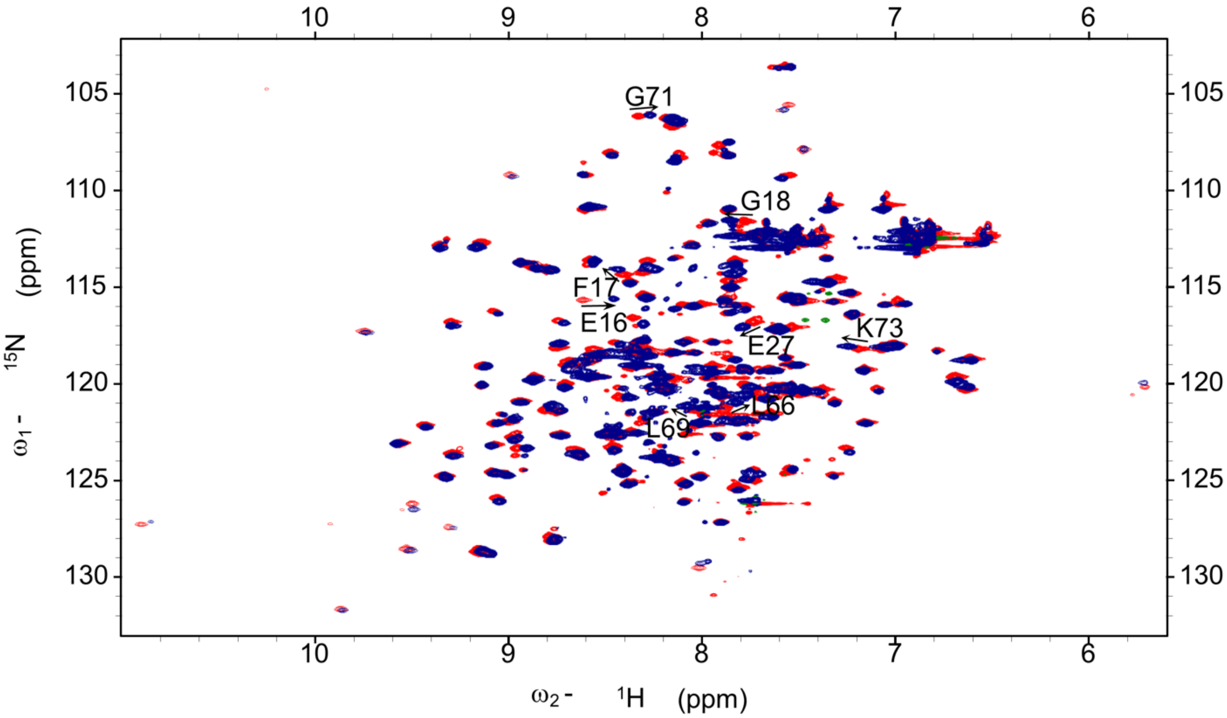


**Figure S1**. $^{1}H$-$^{15}N$ HSQC spectra overlay of ligand-free and IWP-051-bound *Sw* H-NOX. The native *Sw* H-NOX spectrum is colored red and the IWP-051-bound *Sw* H-NOX is colored blue. Residues having significant chemical shift changes are indicated with arrows.

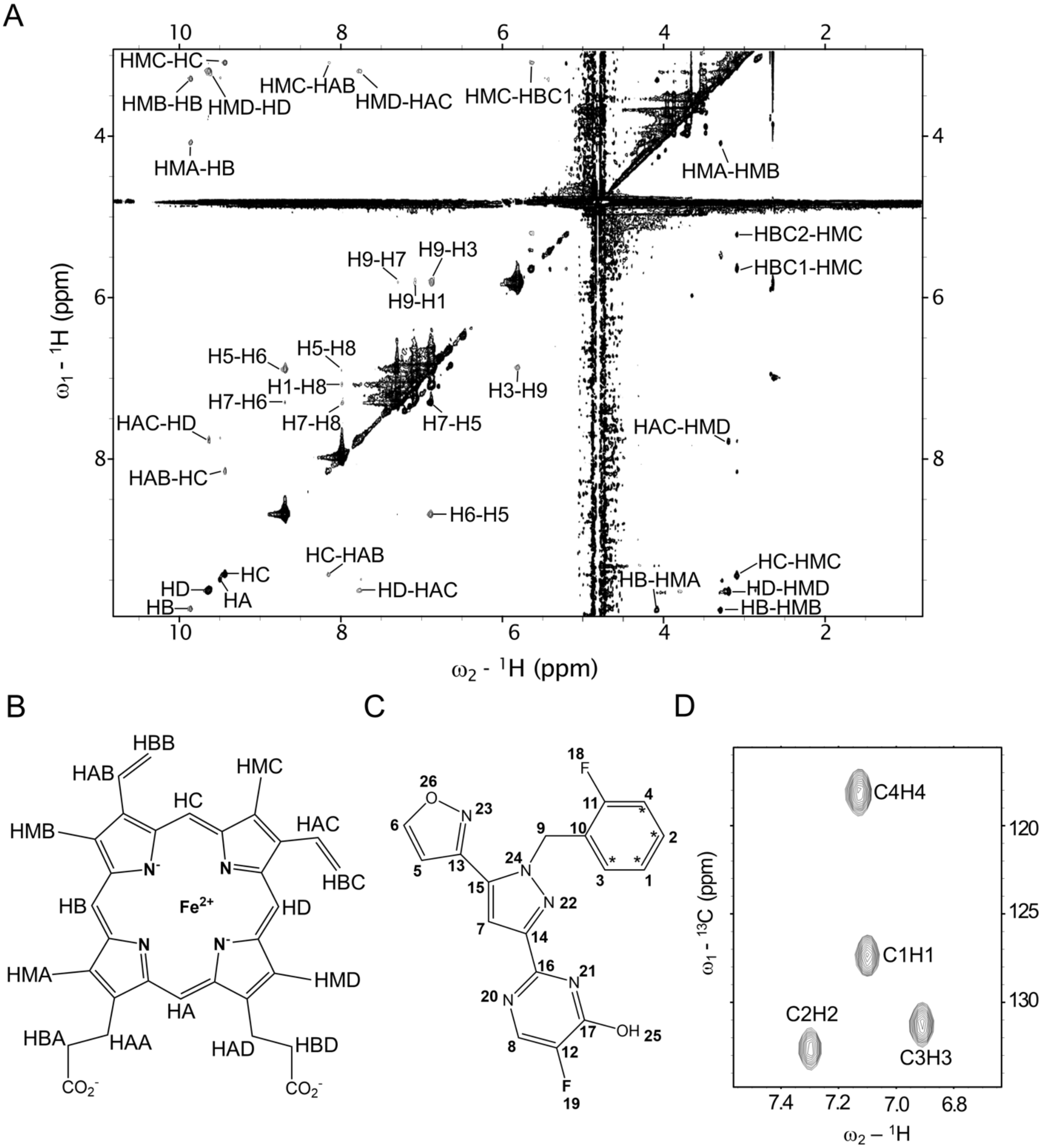


**Figure S2**. Assignments for the protein-bound heme and IWP-051. **A**. 2D [F1,F2]-$^{13}C^{15}N$-filtered NOESY spectrum of the unlabeled heme and IWP-051 bound to $^{13}C$,$^{15}N$-labeled *Sw* H-NOX. Assignments for the intramolecular NOEs are labeled. **B**. Chemical structure of heme with atoms labeled as described in A. **C**. Chemical structure of IWP-051 with atoms numbered as described in A. The location of $^{13}C$-labeled carbons as shown in D is indicated with stars. **D**, 2D $^1H$-$^{13}C$ HSQC spectrum of the 13C-benezyl-labeled IWP-051 bound to unlabeled *Sw* H-NOX. Assignments for the $^1H$ and $^{13}C$ resonances are labeled.

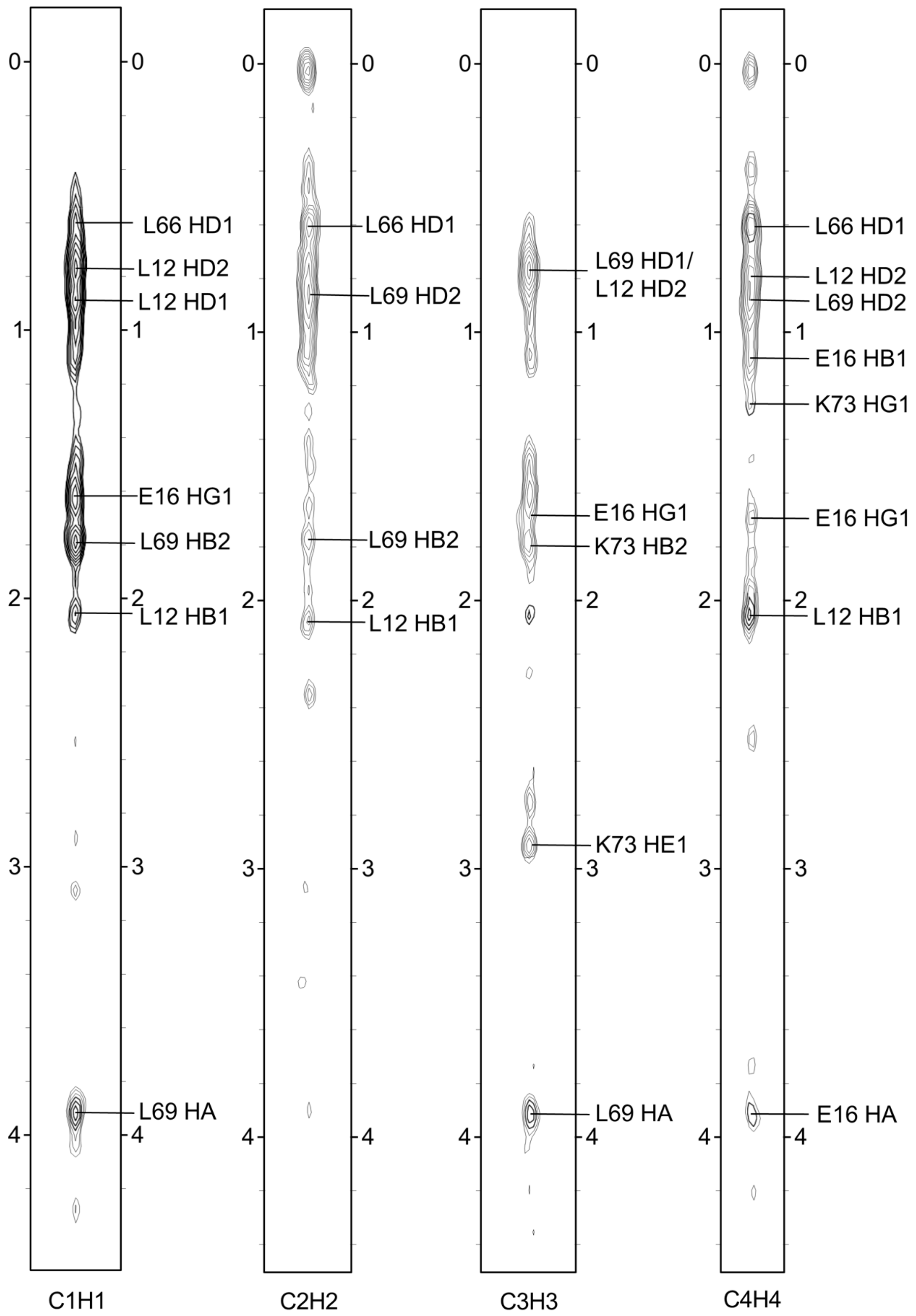


**Figure S3**. Segments from a 3D $^{13}$C-NOESY-HSQC measured on $^{13}$C-benezyl-labeled IWP-051 bound to $^{15}$N-*Sw* H-NOX. Assignments for IWP-051-protein NOEs are labeled.

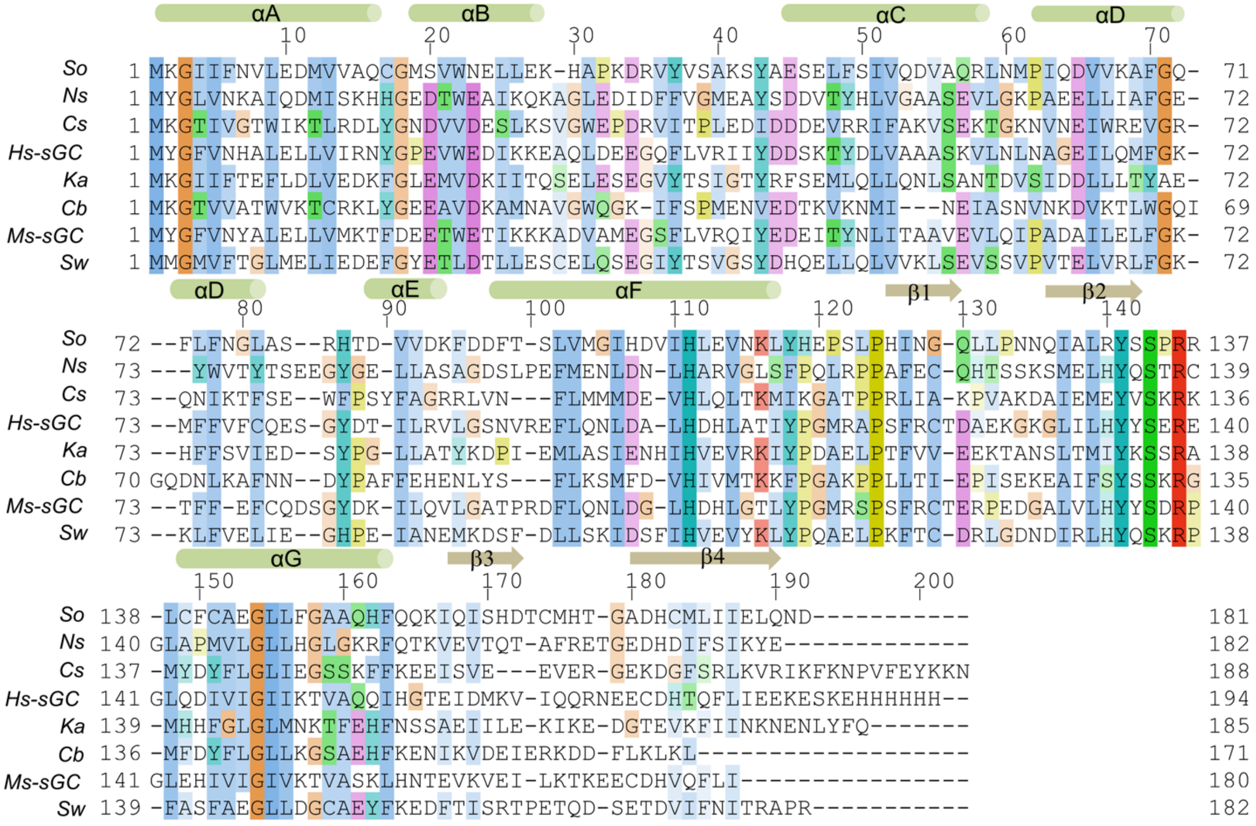


**Figure S4**. Multiple sequence alignment of H-NOX domains. *So*: *Shewanella oneidensis*, *Ns*: *Nostoc sp*, *Cs*: *Caldanaerobacter subterraneus*, *Hs*-sGC: human sGC β H-NOX, *Ka*: *Kordia algicida*, *Cb*: *Clostridium botulinum*, *Ms*-sGC: *Manduca sexta* β H-NOX, Sw: *Shewanella woodyi*. Secondary structure elements are labeled on top based on the structure of *Sw* H-NOX, with α-helices shown as cylinders and β-strands as arrows. The alignment was prepared using the program T-Coffee [38] and the graphical visualization with the program Jalview [39] using the Clustal X color scheme.

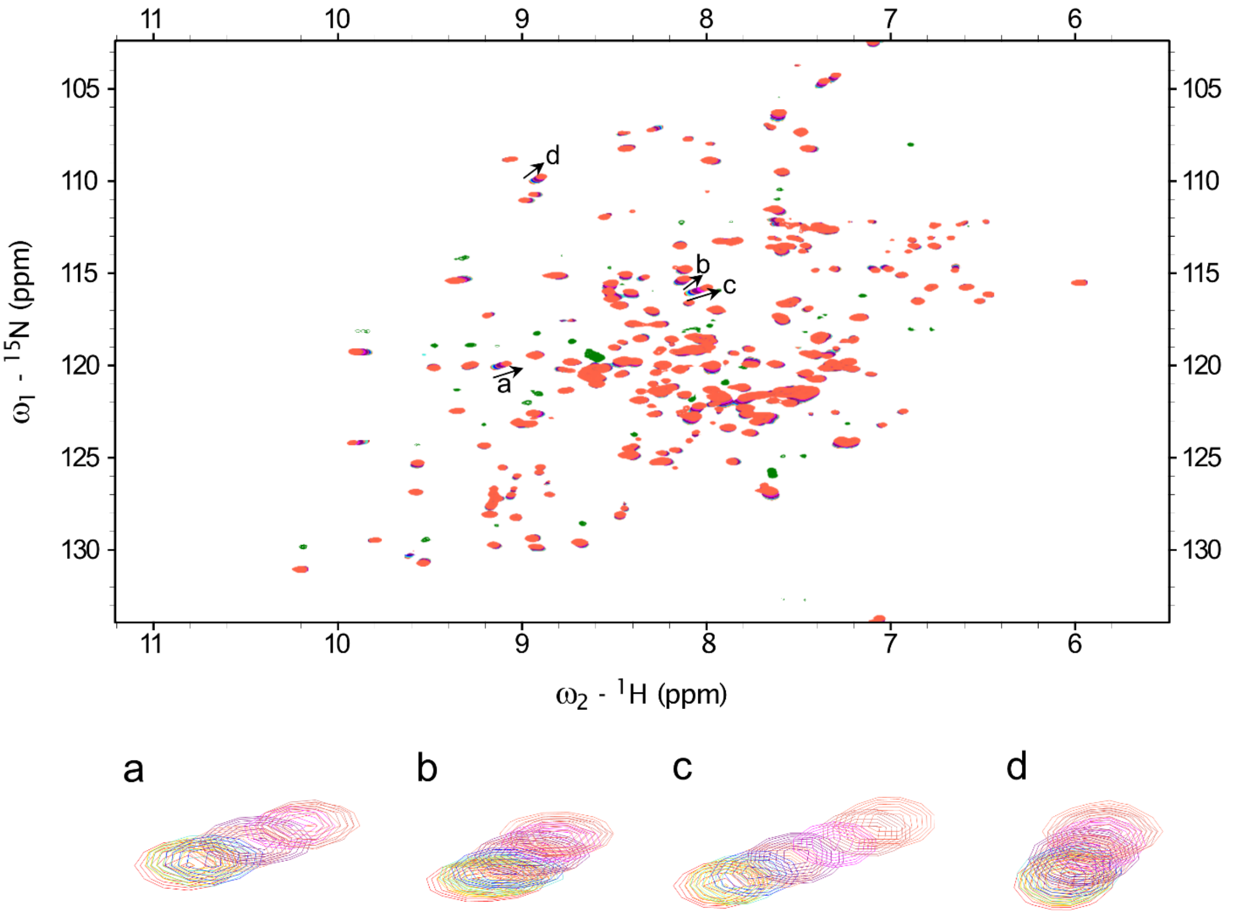


**Figure S5**. Superimposed $^{1}H$-$^{15}N$ HSQC spectra of *Ns* H-NOX titrated with IWP-051. The following ligand/protein ratios are indicated: 0 (red), 0.22 (orange), 0.44 (yellow), 0.88 (turquoise), 1.77 (blue), 2.66 (coral), 3.55 (purple), 5.33 (magenta), 7.11 (indian red), 8.88 (tomato). The lower panel displays an expanded region of residues indicated by arrows in the upper panel.

**Table S1.** Structure statistics

| | *Sw* H-NOX | *Sw* H-NOX-IWP-051 |
|---|---|---|
| Distance restraints | 4434 | 4342 |
| Intraresidue | 1330 | 1327 |
| Sequential $\|i – j\| = 1$ | 972 | 954 |
| Medium-range $1<\|i – j\|<5$ | 1111 | 1084 |
| Long-range $\|i – j\| > 4$ | 1021 | 977 |
| Hydrogen bonds[a] | 192 | 190 |
| Dihedral angle constraints[b] | 249 | 251 |
| Residual dipolar couplings (NH) | 157 | 163 |
| Dipolar coupling R-factor of $D_{NH}$ (%)[c] | 4.77 ± 0.21 | 2.74 ± 0.11 |
| Heme | | |
| Intra-Heme restraints | 15 | 15 |
| Heme-protein restraints | 109 | 113 |
| IWP-051 | | |
| Intra-IWP-051 restraints | N/A | 6 |
| IWP-051-protein restraints | N/A | 24 |
| Ramachandran plot summary[d] (%) (residue 1-29, 45-182) | | |
| Most favored regions | 92.1 | 90.0 |
| Additionally allowed regions | 6.6 | 9.2 |
| Generously allowed regions | 0.7 | 0.2 |
| Disallowed regions | 0.7 | 0.6 |
| Deviations from idealized geometry | | |
| Bonds (Å) | 0.017 | 0.017 |
| Angles (°) | 1.7 | 1.7 |
| Mean pairwise rmsd (residue 1-29, 45-182) | | |
| Backbone (Å) | 0.318 | 0.337 |
| Heavy atoms (Å) | 0.704 | 0.438 |
| Structure quality factor[d], Z-scores | | |
| Procheck G-factor (phi/psi only) | 0.20 | -0.24 |
| Procheck G-factor (all) | -0.77 | -1.24 |
| Verify3D | -6.26 | -6.42 |
| Prosall (-ve) | -0.79 | -0.70 |
| MolProbity clashscore | -0.88 | -1.67 |

[a]Two restraints per hydrogen bond (dHN-O ≤ 2.00 Å and dN-O ≤ 3.00 Å) were implemented for the hydrogen bonds predicted from AUDANA algorithm.

[b]Dihedral angle restraints were generated by TALOS-N on the basis of backbone atom chemical shifts and by analysis of local NOE patterns.

[c]The R factor for residual dipolar coupling is defined as the ratio of the rmsd between observed and calculated values to the expected RMS deviation if the vectors were randomly distributed.

[d]Determined using Protein Structure Validation Suite version 1.5. The suite includes Procheck, Verify3D, Prosall, and MolProbity.
